# Reduced repertoire of cortical microstates and neuronal ensembles in medically-induced loss of consciousness

**DOI:** 10.1101/358168

**Authors:** Michael Wenzel, Shuting Han, Elliot H. Smith, Erik Hoel, Bradley Greger, Paul A. House, Rafael Yuste

## Abstract

Medically-induced loss of consciousness (mLOC) has been linked to a macroscale break-down of brain connectivity, yet the neural microcircuit correlates of mLOC remain unknown. We applied non-linear t-stochastic neighbor embedding (t-SNE) and Lempel-Ziv-Welch complexity analysis to two-photon calcium imaging and local field potential (LFP) measurements of cortical microcircuit activity across anesthetic depth in mice, and to micro-electrode array recordings in human subjects. We find that mLOC disrupts population activity patterns by i) a reduction of discriminable network microstates and ii) a reduction of independent neuronal ensembles. These alterations are not explained by a simple reduction of neuronal activity and reveal abnormal functional microcircuits. Thus, normal neuronal ensemble dynamics could contribute to the emergence of conscious states.

## INTRODUCTION

Medically-induced LOC is generated in millions of patients every year worldwide enabling life-saving surgical procedures or critical care, and is also a hallmark of incompletely understood disorders such as vegetative state (VS). Despite its fundamental importance, the neural circuit and network mechanisms underlying LOC have remained unclear. Several brain areas have been implicated in causing the loss or recovery of consciousness, such as the hypothalamus (Herrera et al., 2016), thalamus (Castaigne et al., 1981), basal ganglia and claustrum (Crick and Koch, 2005; Mhuircheartaigh et al., 2010), or the brain stem (Minert et al., 2017; Moruzzi and Magoun, 1949; Penfield, 1954). Although thalamocortical connections have been central to research on LOC (Flores et al., 2017; Herrera et al., 2016; Penfield, 1954; Steriade, 2003), the role of the cerebral cortex itself remains controversial (Merker, 2007). Early on, animal studies by Lashley (Lashley, 1929) and Pavlov (Pavlov, 1927), or surgical procedures on epilepsy patients (Penfield, 1954) described the cortex’s pivotal role in cognition and learning, yet the removal of even expansive cortical areas resulted in no apparent change of consciousness levels (Lashley, 1929; Pavlov, 1927; Penfield, 1954; Scoville and Milner, 1957). However, diffusely damaged cortex, for example after hypoxia, might contribute to LOC (Jennett, 2002). Recently, fMRI studies investigating large-scale networks have identified a break-down of functional connectivity between cortical macro-areas during mLOC (Barttfeld et al., 2015; Hudetz et al., 2015; Lewis et al., 2012). In addition, one study reported that local network dynamics during mLOC remain similar to those in the conscious state (Lewis et al., 2012), impliying that LOC essentialy arises from discoordination of neural activity across brain areas. Yet, to date, no investigation has employed techniques with sufficient spatial resolution to properly examine the basic neural signatures of LOC at the scale of cortical microcircuits (Tononi et al., 2016).

A general theoretical framework suggests that consciousness depends on the brain’s ability to discriminate between a specific sensory input and a large set of alternatives (Tononi, 2008). In basic agreement, several recent studies have identified a rich set of discriminable resting states at the macroscale of cortical activity (Barttfeld et al., 2015; Hudetz et al., 2015). With such a theoretical framework in mind, along with Nyquist’s foundational work in telegraphy showing that the transmission rate of information logarithmically depends on the set of symbols used (Nyquist, 1924), we hypothesized that an individual’s ability to discriminate between a set of alternatives at any moment should be rooted in discriminable micro-patterns of activity (microstates) at the level of local neuronal ensembles, i.e., the synchronized activation of a group of neurons, which have been postulated to represent functional building blocks of cortical function (Carrillo-Reid et al., 2017; Miller et al., 2014). If this is the case, LOC could arise from alterations in the local microcircuit, which would secondarily generate macroscale connectivity deficits. As a consequence, microstate dynamics across anesthetic depth and recovery could provide important mechanistic insights into the basic building blocks of LOC and this information could help in clinical cases. Here, we provide empirical evidence in mice and humans, across cortical areas, using two different anesthetic agents (isoflurane or propofol) and cellular resolution recording techniques (2-photon microscopy or micro-electrode arrays), that mLOC is indeed associated with a decreased cortical repertoire of discriminable states at the microscale with altered neuronal ensembles. Our results further indicate that local ensemble activity patterns undergo fragmentation during anesthesia. In conjunction with previous macroscale studies, this suggests that during mLOC functional connectivity of the cortex breaks down across spatial scales and that coactive neuronal ensembles are building blocks of cortical function.

## RESULTS

### Reduction of cortical microstates and ensemble fragmentation during mLOC in mice

To explore the role of cortical microcircuits in LOC states, we monitored the activity of cortical populations by combining LFP and fast two-photon calcium imaging (30Hz, 400×400μm field of view) (Yang and Yuste, 2017; Yuste and Denk, 1995; Yuste and Katz, 1991) of head-restrained Thy1-GCaMP6F mice (Dana et al., 2014), allowed to move on a running wheel (Fig. 1A,B). In order to ensure seamless transitions between wakefulness and different anesthetic depths without setup adjustments, a custom tube for the delivery of the inhalatory anesthetic isoflurane was placed in front of the mouse. All animals were accustomed to the microscope, experimental setup, and experimenter before an experiment. In seven mice, we assessed 5 consecutive conditions (10min each) that were matched across animals based on hallmark LFP patterns (Fig. 1C): wakefulness, light sedation (~0.5% isoflurane partial pressure in air, slowed LFP yet similar to wakefulness), surgical anesthesia (~1.0% iso, continuously increased LFP delta [1-4Hz] spectral power), deep anesthesia (~1.5% iso, burst-suppression LFP), and recovery (iso off). In order to further index the animals‘ level of consciousness, we assessed clinical parameters (breath-rate, reaction to tail pinching), and locomotion recorded by an infra-red sensor at the wheel (Fig. 1D). During surgical or burst-suppression anesthesia, animals were also unresponsive to tail-pinching, while after halting isoflurane, they remained so initially during the wake-up period. Calcium transients of registered neurons were extracted from image series (Fig. 1B, E), pre-processed and binarized (Fig. 1F, see methods). To group together similar patterns of population activity from the binarized spike raster matrices, we performed t-distributed stochastic neighbor embedding (t-SNE (van der Maaten, 2008), Fig. 1G, please see methods) leading to a 2dimensional embedding space from which clusters could be identified by watershed segmentation (Suppl. Fig. 1A). Identified clusters were confirmed by comparing intra-versus inter-cluster distances (Suppl. Fig. 1B). Across 7 mice, a total number of 851 neurons (122 ± 12 s.e.m.), and 34,321 active frames (4,903 ± 929 s.e.m.) participating in 425 microstates (61 ± 9 s.e.m.) were analyzed. Across all mice, locomotion was strongly reduced during mild anesthesia and recovery, and no locomotion was detected during surgical or burstsuppression anesthesia (Fig. 2A).

**Figure 1.**
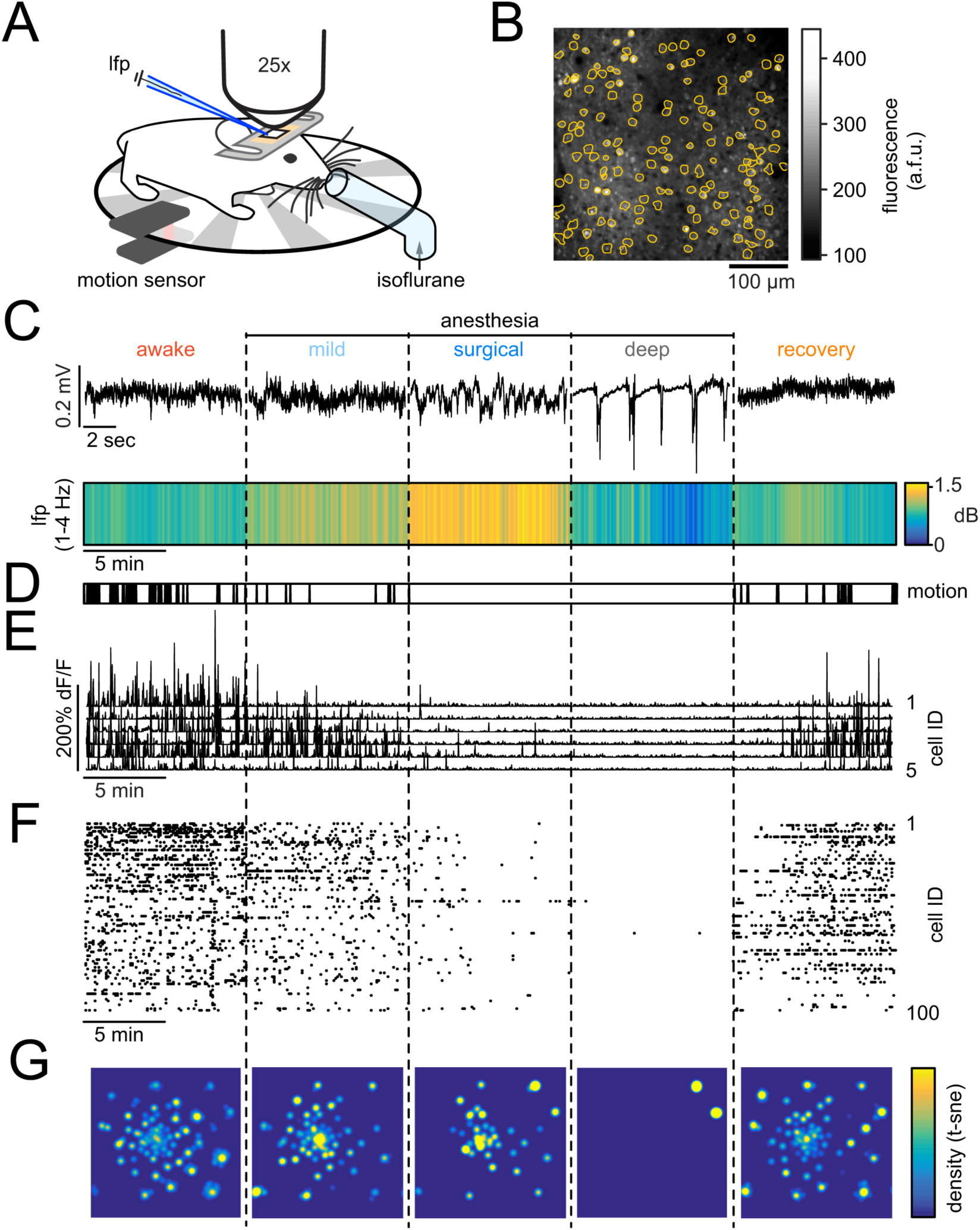
Monitoring microcircuit signatures of mLOC in mice. **A)** Awake, head-restrained mouse on a running wheel. Movement was measured by an infrared sensor. For seamless transitions across conditions, isoflurane was delivered through a custom tube placed right in front of the mouse. For LFP recordings, a pulled glass microelectrode was carefully inserted into the cortex at around 250μm depth through a small burr hole next to an implanted glass cover. Through the glass, two-photon calcium imaging was performed. **B)** Image of typical field of view and registered neuronal somata outlines (orange). **C)** Upper panel: Representative brief raw LFP traces across five conditions. Lower panel: Avg LFP delta range [1–4Hz] spectral power across all five 10min long conditions (n=7 animals). Note the continuously increased delta power during surgical anesthesia. **D)** Superimposed motion of all 7 mice across conditions. Note that movement is absent in surgical and burst-suppression anesthesia. **E)** Calcium transients of 5 representative registered neurons across all five conditions. **F)** Corresponding raster plot of all registered neurons. **G)** Density map of microcircuit states, visualized by t-SNE for the entire experiment, displayed per condition. Vectors representing the population activity at each time point were transformed into a two-dimensional space while preserving local structures, and a density map was generated from the scatter plot.

**Figure 2.**
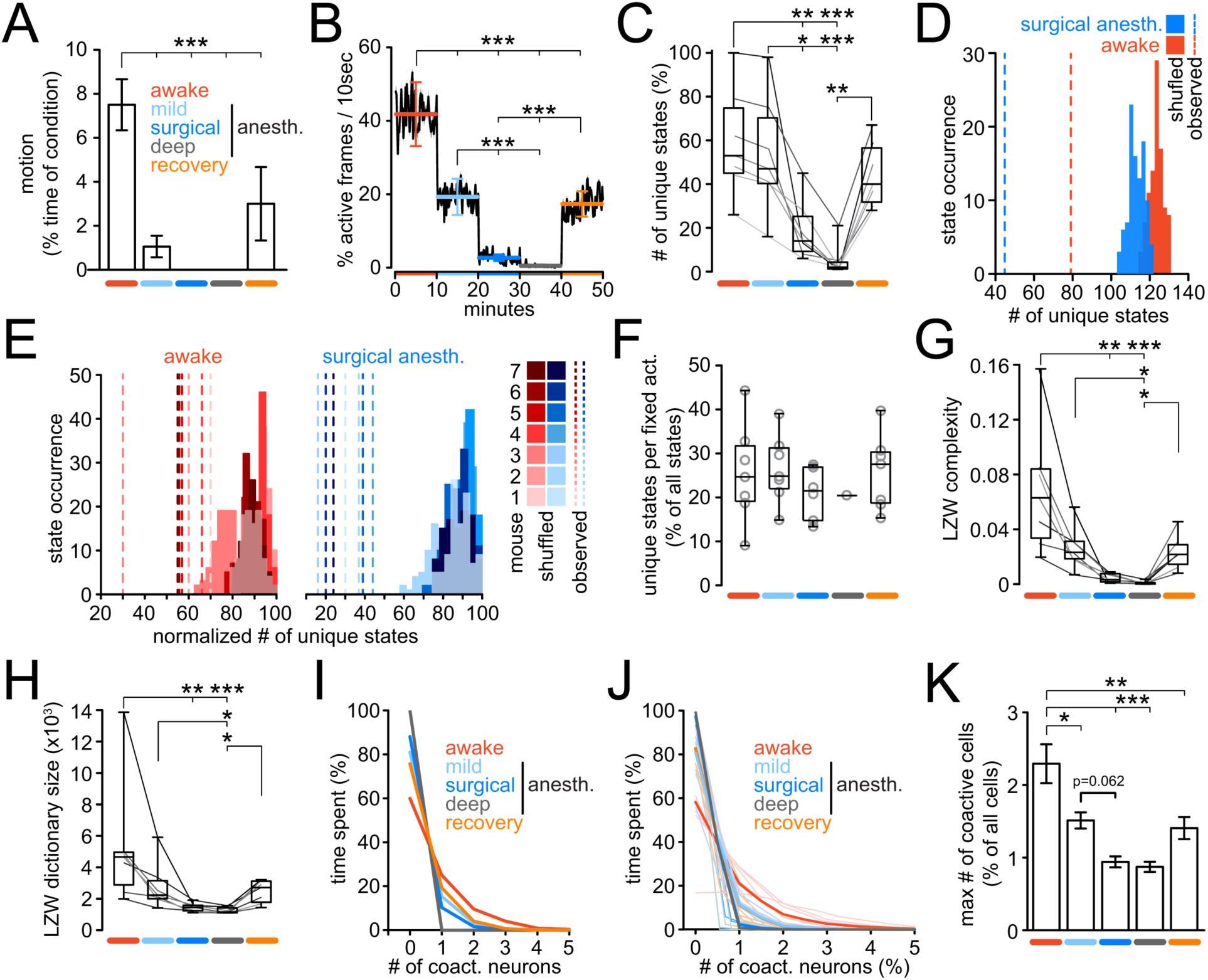
Reduced repertoire of cortical microstates and neuronal ensembles upon mLOC in mice. **A)** Quantification of locomotion across the entirety of each experiment (exp.), displayed as % locomotion per total 10min long condition; anesth.=anesthesia. **B)** Neural activity level across conditions, quantified as probability of detecting activity within a moving 10sec window in a yes or no fashion; errorbars represent means ± s.e.m. per condition; conditions colored as in a). **C)** Boxplots of number of unique microstates across conditions as % of all identified unique microstates in a given experiment; line plots are individual experiment (grayscale) **D)** Representative experiment.: total number of observed unique microstates (dashed lines) during wakefulness (red) or surgical anesth. (blue) vs. corresponding distributions of values from 100 randomized datasets. No overlap of observed vs. random data (p<0.01). **E)** Same comparison as in d), across all exp. each max-normalized for purpose of visualization. In all exp. no overlap of observed vs. random data, equal to p<0.01. **F)** Boxplots of number of unique microstates visited across specific conditions per fixed overall activity (any window of 50 accumulative active events in raster matrix), displayed as % of all states per exp.. **G)** Boxplots of LZW complexity across conditions; line plots are individual exp. (grayscale) **H)** Boxplots of LZW dictionary size across conditions; line plots are individual exp. (grayscale) **I)** Representative exp.: Number of co-active neurons versus the time spent by population in frames containing such co-activity, displayed per condition. **J)** Same comparison as in **I)**, containing all exp..; number of coactive neurons as % of all neurons per exp. for purpose of visualization. Thick lines represent means, thin lines individual exp. conditions **K)** Maximum number of co-active neurons (as % of all neurons per exp.) participating in a specific microstate, across conditions (colors as in a)). Borderline statistical significance mild vs. surgical anesthesia p=0.062. Except for representative examples in D) and I), all data in figure 2 from n=7 animals; all errorbars represent mean ± s.e.m.; all boxes in boxplots represent 25–75%ile of the data, bars within boxes represent means. Except for comparisons between observed and randomized data, all statistical analyses represent 1way-anova with Bonferroni post-test. *p<0.05, **p<0.01, ***p<0.001.

Consistent with previous studies, neural firing dropped promptly during mLOC (Ishizawa et al., 2016; Lewis et al., 2012), with total neuronal activity during surgical anesthesia falling below 10% of baseline (Fig. 2B). Intriguingly, and in contrast to Lewis et al. (Lewis et al., 2012), the number of discriminable microstates, that is, the number of identifiable watershed regions in the embedding space, was strongly reduced during surgical and burst-suppression anesthesia (Fig. 2C). However, despite significantly reduced overall neural activity, most microstates found during wakefulness were still present during mild anesthesia and again during recovery, (Fig. 2B). To control for the potential influence of the number of coactive neurons on the microstate classification during each anesthetic condition, we repeated the analysis shown in Figure 2D but only using frames during the awake condition that contained no more than the maximum number of coactive cells observed at least once during surgical anesthesia (2.6 ± 0.4 s.e.m., n=7 mice), with consistent results (Suppl. Fig. 2A). In other words, experimental conditions matched to have the same maximum number of active cells had fewer microstates in data from anesthetic conditions vs. wakefulness. To explore if the effect of microstate reduction during deeper anesthesia could be explained by the random coactivation of neurons, we repeated t-SNE and watershed segmentation on 100 randomized datasets derived from observed data through within-frame shuffling. This procedure preserved total per-frame-activity while disrupting within-frame patterns (Suppl. Fig. 1C,D). Importantly, in all animals and experimental conditions, the number of randomly generated microstates was significantly higher than that found in the corresponding observed data (Fig. 2D,E; shown is awake and surgical anesthesia. See also suppl. Fig. 1D), except for the wake-up period in one experiment and burst-suppression anesthesia. This lack of significant difference could be explained by a critically low overall activity. These results indicated i) that the observed microstates were non-random, given sufficient activity, and represented a much smaller set compared to all possible states, and ii) despite reduced activity during surgical anesthesia, a large number of discriminable microstates would still be theoretically possible. Interestingly, when we normalized the microstates by the neuronal activity, by measuring the number of unique microstates in any window accumulatively containing 50 events of neural activity, we found that across conditions (for burst-suppression in one animal), the local population cycled through a similar number of unique microstates per fixed activity (Fig. 2F). Due to this result, we sought to investigate, whether the reduction of microstates during deeper anesthetic conditions could be explained simply by a sparser appearance of individal microstates as a consequence of the much reduced activity, and not by a true reduction of the number of microstates occuring during anesthesia. To this end, we performed experiments to compare the total number of unique microstates over 10 minutes during wakefulness against the number of unique microstates occuring over 50 minutes of surgical anesthesia. Even over this much extended period of recording, with the corresponding increase in total neuronal activity, the number of microstates remained strongly reduced (Suppl. Fig. 2B,C). Together, these results indicated that the local network indeed drew from a reduced repertoire of discriminable microstates during anesthesia, yet cycled through a similar set of unique states given sufficient overall activity.

To minimize the possibility that our results could be due to our dimensionality reduction and clustering approach, we applied Lempel-Ziv-Welch compression (LZW, suppl. Fig. 2D) as a complementary analysis based on different algorithmic principles (Welch, 1984; Ziv and Lempel, 1978). In accordance with our t-SNE results, the Lempel-Ziv-Welch complexity (LZWC) was increasingly lower across anesthetic depth (Fig. 2G). Furthermore, and corroborating the number of microstates identified by t-SNE and watershed segmentation, the number of unique features (’words’, or collectively, ’dictionary’) encountered by LZW in a spike matrix also decreased significantly across anesthetic depth (Fig. 2H).

While performing the experiments for this study, we became aware that changes in ensemble co-activity seemed to be associated with mLOC. To study this, we operationally defined ensemble as a group of neurons that is coactive in any given frame (Miller et al., 2014). Indeed, neural co-activity within a specific microstate consistently decreased with anesthetic depth (Fig. 2I-K). These neuronal ensembles were also not due to chance firing of cells, as within-cell shuffling, disrupting co-activity patterns while maintaining the same total activity per condition (Suppl. Fig 1C, lower panel), led to random distributions significantly smaller than the observed data (Suppl. Fig 3A,B). These results indicated a reduction of local ensembles (Miller et al., 2014) during mLOC, with a progression towards independent single neuron activity patterns during deeper anesthesia.

### Reduction of cortical microstates and ensemble fragmentation during mLOC in humans

To investigate whether our mouse findings held true in human cortex, we sought to study discriminable microstates and neural coactivity at fine anatomical scale across anesthetic depth in two neurosurgical patients. These were epilepsy patients undergoing anesthesia induction for an epileptic focus resection following their clinical monitoring. In addition to subdural macro-electrode implantation for epileptic focus identification in the temporal lobe (Fig. 3A), these patients were implanted with a microelectrode-array (4 × 4mm, 96 electrodes, Fig. 3A,B) capable of measuring both LFP (Fig. 3C), and single unit activity (SUA) in the anterior middle temporal gyrus (Fig. 3D). In both patients, three anesthetic conditions (mild, surgical, and burst-suppression) were examined by applying intravenous propofol boluses. Anesthetic conditions were determined based on LFP (Fig. 3C), as well as the bispectral index score, a frequently used intraoperative index of anesthetic depth. Since the micro-electrode array was implanted on the day clinical electrodes were removed, neither wakefulness nor the wake-up period could be recorded. As with the mouse analysis, neural firing was binarized (Fig. 3D), and similar activity patterns across time were first identified using t-SNE, and watershed cluster segmentation (Fig. 3E, suppl. Fig. 1A). In the two human subjects, a total of number of 145 single units (72.5 ± 18.5 s.e.m.), and 6,950 active frames (3,475 ± 1147 s.e.m., 1 frame = 100ms epoch) participating in 89 microstates (44.5 ± 5.5 s.e.m.) were identified. As in mice, a general drop of neural activity was observed with increasing anesthetic depth (Fig. 3F). Likewise, our finding of a decrease of the number of discriminable microstates during anesthesia in mice held true in both human patients (Fig 3G). As before, the effects could not only be explained by a random coactivation of neurons, as observed results were significantly different from randomized datasets (Fig 3J). In addition, LZWC and LZW dictionary size both decreased with anesthetic depth, too (Fig 3H-J). Thus, in line with our results in mice, the local population drew from an increasingly reduced repertoire of microstates with increasing anesthetic depth. Finally, in both patients, neural coactivity patterns non-randomly disappeared with increasing depth of anesthesia (Fig 3K-L, suppl. Fig. 3C).

**Figure 3.**
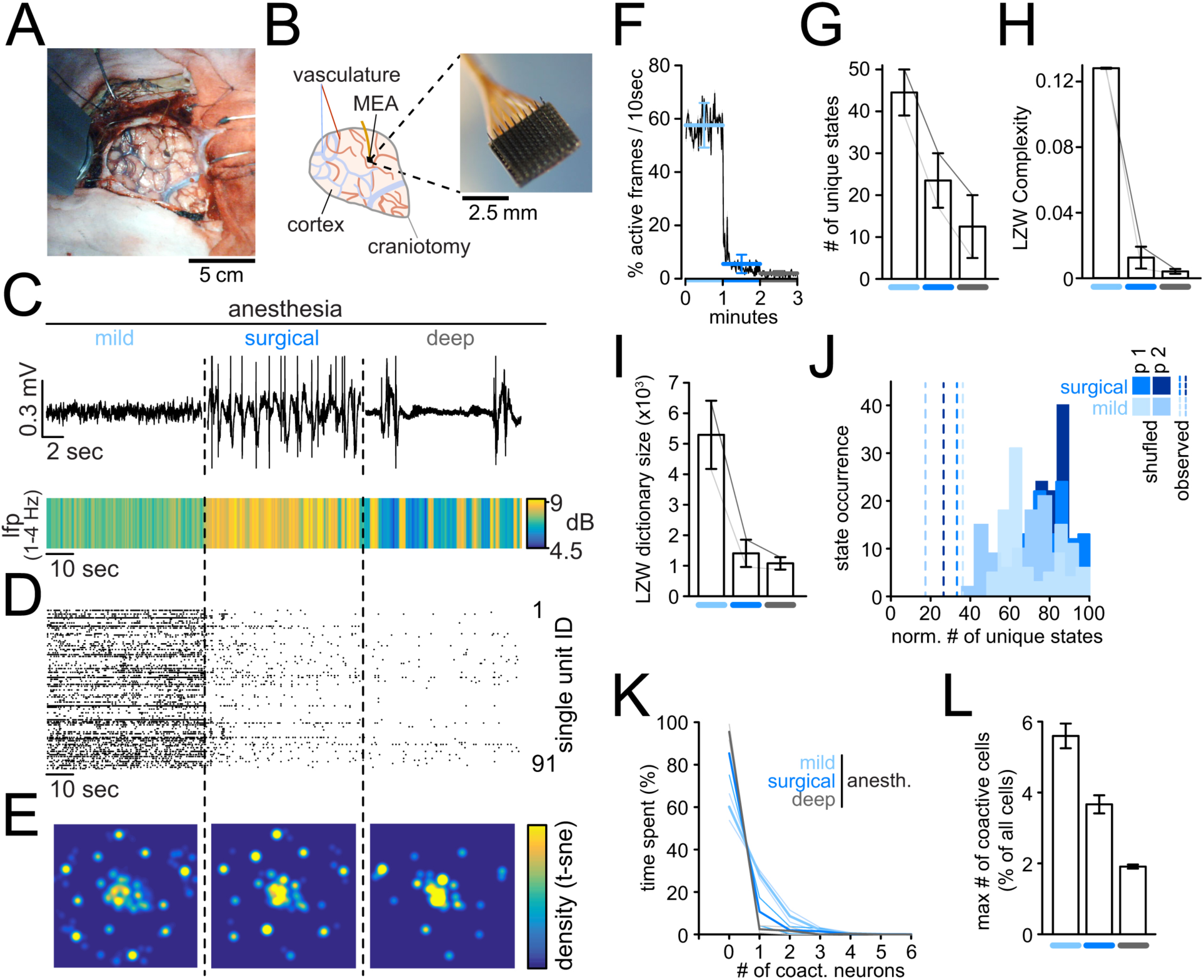
Reduced repertoire of cortical microstates and ensembles upon mLOC in humans. All errorbars in figure 3 represent mean ± s.e.m.. (n=2 patients). **A)** Photo of the craniotomy carried out on one patient in this study. **B)** Left: Drawing of craniotomy in a). craniotomy (gray); arteries (red); veins (blue), multi-electrode microarray (MEA, black). Right: close-up photo of micro-electrode array (4×4mm, 96 electrodes). **C)** Upper panel: Representative brief raw LFP traces across three anesthetic conditions in one patient. Lower panel: Avg LFP delta range [1–4Hz] spectral power across conditions, and patients. **D)** Raster plot of all single units in one patient. **E)** 2-dimensional density plot of same data after t-SNE, displayed per condition. Every dot represents a timepoint containing neural activity. **F)** Neural activity level across conditions, quantified as probability of detecting activity within a moving 10sec window in a yes or no fashion; conditions colored as in c). **G)** Number of unique microstates across conditions; line plots are individual patients (grayscale). **H)** LZW complexity across conditions; line plots are individual patients. **I)** LZW dictionary size across conditions; line plots are individual patients. **J)** Number of observed unique microstates (dashed lines) during mild anesth. (light blue) or surgical anesth. (dark blue) vs. corresponding distributions of values from 100 randomized datasets, max-normalized for purpose of visualization. No overlap of observed vs. random data, equal to p<0.01; p1/2 represent patient 1/2 **K)** Number of co-active neurons versus the time spent by population in time points containing such co-activity per condition, displayed as mean across patients (thick lines). Thin lines represent conditions in individual patients. **L)** Maximum number of co-active neurons (as % of all neurons per exp.) participating in a specific microstate across conditions.

## DISCUSSION

In this study, we used single cell resolution population recording techniques in mice and human subjects to investigate changes in microscale activity patterns during wakefulness, across anesthetic depth, and recovery from anesthesia. In contrast to recent publications (Hudetz et al., 2016; Lewis et al., 2012), we find that, across species, cortical areas and anesthetic agents, mLOC is associated with a decreased cortical repertoire of discriminable microcircuit states and neuronal ensembles. Moreover, that the number of microstates found in all conditions represent a small fraction from all possible microstates, indicating that these microstates are not due to chance coactiavtion of neurons. To ensure that the observed phenomenon was robust across analytical approaches, we applied two analyses that are based on different algorithmic principles (t-SNE and LZW). In addition, as these methods have not yet become commonplace in studying two-photon calcium population imaging or MEA spike raster matrices, we finally used a simple PCA approach as a third and widely used method, with consistent results (Suppl. Fig. 2 E,F).

In both mice and humans, we find a decrease in neuronal activity during mLOC, as it has been reported before. But our main finding is that the number of functional microstates and ensembles is selectively reduced during mLOC. Certainly, the decrease of overall activity upon start of anesthesia contributes to the reduced number of microstates observed here during deeper anesthesic conditions. And, as the reducion of general activity constitutes an inherent feature of general anesthesia, it is practically impossible to completely account for this effect. However, even when we record local network activity under surgical anesthesia for five times longer than during wakefulness, we find a reduced number of discriminable microstates (Suppl. Fig. 2B,C). This means that the reduced number of states or ensembles cannot be simply accounted for by the reduced activity levels and represents a specific malfunction of the microcircuit. Indeed, a drop of general activity does not inherently result in a reduced number of discriminable states. This notion is supported by simulated spiking data, in which 10 neurons independently fired once (1 cycle) or ten times (10 cycles) over the course of 100 frames (Suppl. Fig. 2G). As a result, one matrix contained ten times less overall activity than the other, yet the number of microstates that were identified by t-SNE, remained exactly the same. This underscores that an equivalent number of microstates observed during wakefulness would technically be possible during deeper anesthesia, too. However, our data indicate that the reduced number of discriminable microstates and ensembles during deeper anesthetic conditions is indeed due to a reduced repertoire available to the local network.

Further, we find that local ensemble activity patterns undergo fragmentation during mLOC, that is, coactivity patterns disappear with increasing anesthetic depth. The discrepancy between our findings and previous literature might be explained by the different recording techniques used, and how the obtained signals were analyzed. Using a 64-electrode microarray, Hudetz and et al. described that the repertoire of mesoscale cortical activity is not altered during anesthesia. However, their conclusion was based on the analysis of local field potentials filtered at 4-60 Hz, which means that they did not detect local neural firing. This could explain the mismatch between their finding and ours, because neural input dynamics and actual neural firing can often differ. Using a 96-electrode microarray similar to the one used in our study, Lewis and colleagues reported that while macroscale connectivity broke down during mLOC, microscale activity would persist similarly to the conscious state, yet in an insular manner. Of important note, this conclusion related to the persistence of microscale activity was based on a pairwise correlation analysis of only a small fraction (<15%) of recorded single units showing the highest firing rate in one human individual. It remains debatable whether this pre-selection and pairwise correlation analysis was the optimal analytical approach, given that a large body of work has shown generally sparse neural firing in the cortex (reviewed in (Barth and Poulet, 2012)), and weak pairwise correlations between local neurons in many cortical areas, under various circumstances (e.g. reviewed in (Cohen and Kohn, 2011) containing a table of 26 studies in primates with a mean pairwise neural correlation coefficient of 0.107). In the context of both aforementioned reports, it is further important to keep in mind that weak pairwise correlations of local neural firing may neither preclude synchronous network states (Schneidman et al., 2006), nor strong computational properties of the network (Hopfield, 1982). In our study, all units belonging to the recorded local network, in two patients, were included in the frame by frame analysis using t-SNE or LZWC across anesthetic depth, which in our view more accurately covers changes of microscale network activity patterns. Finally, the results in humans were consistent with our findings in mice using cellular resolution two-photon calcium population imaging.

To conclude, extending previous studies at the macroscale (Barttfeld et al., 2015; Hudetz et al., 2015; Lewis et al., 2012), our results provide evidence that, during mLOC, the functional connectivity of the cortex also breaks down at the microscale. In fact, as local ensembles undergo fragmentation, functional connectivity and integration of information at the macroscale should naturally be altered. In sum, our study provides a unifying framework for functional brain connectivity changes during mLOC across spatial scales, adding a micro-anatomical basis for how the cerebral cortex, along with subcortical areas in a distributed complex cerebral system, could contribute to loss or re-gain of consciousness (Koch, 2012).

## Methods

### Experimental Model and Subject Details

#### Animal subjects

Experiments were conducted with care and accordance with the Columbia University institutional animal care guidelines. Experiments were carried out on Thy1-GCaMP6F (Dana et al., 2014) adult transgenic mice at postnatal age of 4-6 months. No animals were used for previous or subsequent experimentation. Food and water was provided ad libitum. All mice were kept at a 12 hour light/dark cycle.

#### Human subjects, and Ethics Statement

Two human subjects were included in this study. Patient 1 was a 31-year-old male and patient 2 was a 64-year-old male. Both patients were undergoing neurosurgical resection of the anterior temporal lobe (patient 1: left; patient 2: right) in order to treat medically refractory mesial temporal lobe epilepsy. The University of Utah Institutional review board approved these experiments and both subjects provided informed consent prior to participating in the study.

### Method Details

#### Animals, surgical procedures, and setup acclimatization

Prior to the actual experiment, mice were anesthetized with isoflurane (initial dose 2-3% partial pressure in air, then reduction to 1-1.5%). Right before surgery, all mice received carprofen (s.c.), enrofloxacin (s.c.), and dexamethasone (i.m.). Under sterile conditions, a small flap of skin above the skull was removed and a titanium head plate with a central foramen (7×7mm) was attached to the skull with dental cement above the left hemisphere. A small cranial aperture (around 2×2mm) was established above left somatosensory (coordinates from bregma: x 2,5mm, y −0,24mm, z −0,2mm) or visual cortex (x 2,5mm, y −0,02mm, z −0,2mm) using a dental drill. Then, the craniotomy was covered with a thin glass cover slip (3×3mm), which was fixed in place with a slim meniscus of silicon around the edge of the glass cover and finally cemented on the skull using small amounts of dental cement around the edge. A small section of skull (1×1mm) was left blank next to the cemented glass cover. Postoperatively, all mice received carprofen daily for 3 days. Over the following days, mice were accustomed to the experimenter, and the experimental setup until no signs of distress were present. Mice usually became rapidly acclimatized to the microscope and running on a wheel under head-restrained conditions over the course of 2-3 acclimatization sessions lasting 30 minutes each. Around two weeks after the implant of the glass cover slip, on the day of the actual experiment, mice underwent brief surgery again. Using isoflurane as described above, a small burr hole was established in the area that had been left blank next to the cemented glass cover for access by a glass micropipette for LFP measurements (find a more detailed description under electrophysiology, below). A reference electrode was place over the right frontal cortex. Thereafter, mice were transferred to the microscope for the experiment.

#### Experimental timeline in mice

In each experiment, animals were kept in head-restrained conditions yet allowed to move freely on the running wheel. Throughout each experiment, in addition to local population imaging, cortical activity was recorded by LFP measurements serving as an additional proxy of anesthetic depth aside from clinical assessment (breath rate, locomotion, responsiveness to tail pinching). After the first image and LFP series during wakefulness, mild anesthesia was established by delivery of low concentrations of isoflurane through a plastic cylinder positioned right in front of the mouse’s nose (Fig. 1 A). During mild anesthesia/low levels of isoflurane (0,5% partial pressure in air – ‘ppa’), mice remained responsive to tail pinching. This responsiveness ceased completely once general anesthesia was achieved by increasing the dose of isoflurane to around 1,0% ppa. Then, burst suppression anesthesia was induced through another increase of isoflurane to around 1.5%, and maintained for 10 minutes. Finally, isoflurane delivery was halted, and imaging and lfp recordings continued while the animal was allowed to fully recover from anesthesia. Once the experiment was completed, animals were deeply anesthetized and sacrificed humanely.

#### Infrared locomotion detection in mice

Locomotion was detected using an infrared sensor at the running wheel. The initially transparent running wheel was adapted to block the infrared light from passing through the wheel by using equally spaced strips of light absorbent tape (Fig. 1A). Thus, whenever the mouse would locomote, the light path between the light source underneath the wheel and the sensor on top of it would alternatingly get blocked or released rapidly. During each such transition (the longer the locomotion, the more transitions), the infrared sensor produced a large positive (from blocked to transparent) or negative (transparent to blocked) change in voltage that could be recorded at 1 kHz temporal resolution alongside the LFP using Prairie View Voltage Recording Software. Transitions could be easily extracted after an experiment to create a binary vector of locomotion or rest. For each event in the binary vector, 1 second of locomotion was counted, and the relative time of locomotion versus resting, for each experimental condition, was calculated.

#### Two-photon calcium imaging in mice

Neural population activity in cortical layer II/III was recorded by imaging changes of fluorescence with a two-photon microscope (Bruker; Billerica, MA) and a Ti:Sapphire laser (Chameleon Ultra II; Coherent) at 940 nm through a 25x objective (Olympus, water immersion, N.A. 1.05). Resonant galvanometer scanning and image acquisition (frame rate 30.206 fps, 512 × 512pixels, 150–250µm beneath the pial surface) were controlled by Prairie View Imaging software. Multiple datasets were acquired consecutively over the course of an experiment (in total 100,000–150,000 frames) with several momentary breaks interspersed for reasons of technical practicality.

#### Two-photon image processing

Active cells were first identified visually in raw tiff movie files using ImageJ software (Schneider et al., 2012). 10 minutes of imaging during each of the five anesthetic conditions (matched across animals by raw LFP, spectral power, and clinical parameters) were concatenated into one large movie tiff file spanning the entire experiment. A list of cell centroid spatial coordinates was obtained and used to initialize the recently described constrained nonnegative matrix factorization algorithm (CNMF) to extract calcium transients of all registered cells in MATLAB (Pnevmatikakis et al., 2016; Yang et al., 2016). Prior to the initialization, tiff series were down sampled (averaged) from the original 30Hz temporal imaging resolution to 10Hz, and 512×512 pixel spatial resolution to 256×256 pixels. The CNMF algorithm finds spatiotemporal components based on pixels of high covariance around defined cell centroids while accounting for background fluorescence and minimizing signal noise. Based on the extracted fluorescence traces, ΔF/F signals are calculated for each cell using a sliding window (30 seconds). To derive binarized activity events from the ΔF/F signals, the ΔF/F is temporally deconvolved with the CNMF parameterized fluorescence decay. In addition, a temporal first derivative (slope) is independently obtained from the ΔF/F signals of individual cells. Then, the deconvolved signal and the derivative are thresholded at at least four standard deviations from the mean signal, respectively. At each time point, if both the devoncolved signal and first derivative exceed the threshold, a binary activity event is detected. The binary matrices obtained in this way, contained the recorded number of neurons across 30,000 frames of imaging (50 minutes), and represented the input matrices for t-SNE embedding and watershed segmentation, or Lempel-Ziv complexity analysis, as described below.

#### Local field potential recordings in mice

For LFP measurements, a sharp glass micropipette (2-5 MΩ) containing a silver chloride wire, back-filled with saline, was diagonally advanced into the cortex (30° angle) under visual control. The pipette was lowered through a burr hole next to the glass cover slip, as described above, to a depth of around 200 µm beneath the pial surface. A reference electrode was positioned over the contralateral frontal cortex. LFP signals were amplified by use of a Multiclamp 700B amplifier (Axon Instruments, Sunnyvale, CA), low-pass filtered (300Hz, Multiclamp 700B commander software, Axon Instruments), digitized at 1 kHz (Bruker) and recorded using Prairie View Voltage Recording Software along with calcium imaging.

#### Single Unit Activity and LFP data acquisition and pre-processing in humans

Human electrophysiological data were acquired from a Utah-style microelectrode array implanted in each subject’s middle temporal gyrus, approximately 3 cm from the temporal pole. Detailed surgical methods are described in House et al (House et al., 2006). Data from these microelectrodes were acquired at 30,000 samples per second and pseudo-differentially amplified by 10 using an FDA-approved neural signal processing system (Blackrock Microsystems, Salt Lake City, UT). Continuous recordings were acquired from the microelectrode arrays while anesthesia was maintained at different anesthetic depths. Signals from the microelectrode arrays were segregated into two data streams: single unit activity (SUA), and local field potentials (LFP). SUA was acquired by first band-pass filtering the voltages recorded on each microelectrode between 300 and 3,000 Hz (4th order butterworth filter). This high pass filtered signal was thresholded at −3.5 times its root mean square, and 48 samples around each threshold crossing were retained for spike sorting. Spike sorting was carried out in a semi-supervised fashion on a feature space of the first three principal components, and clustering using the T-distributed expectation maximization algorithm (Shoham et al., 2003). The time stamps of well-isolated single units were retained for further analysis in a binary matrix in which one dimension represented the activity of a single unit and the other dimension represented time at 1000 samples per second. LFP data were acquired by non-causal low pass filtering (500 Hz fir filter) the voltage recorded on each microelectrode and averaging across channels. This mean LFP across the array was then down-sampled to 1000 samples per second.

#### Bispectral Index Scale in human subjects

Anesthesia depth was measured using a bispectral index scale (BIS) using a BIS Vista monitoring system (Aspect Medical Systems, Norwood, MA). Microelectrode recordings began after both patients were induced at a light anesthesia depth (BIS ≈ 50) with propofol and remifentanil. Propofol anesthesia was then delivered in three discrete intravenous boluses in order to achieve deep anesthesia (BIS ≈ 20).

#### Analysis of LFP spectral power in mice or human subjects

LFP low frequency spectral power (1-4Hz) was calculated with a 1 Hz temporal resolution. A Fast-Fourier transform (FFT) was carried out, and the spectral power was calculated as the squared absolute value of the complex output of the FFT. Finally, spectral power was averaged across the low frequency range.

#### T-SNE and watershed segmentation of mouse or human neural activity

We identified cortical microstates of the neuronal population using an un-supervised nonlinear embedding method, t-Distributed Stochastic Neighbor Embedding (t-SNE) (van der Maaten, 2008). T-SNE was performed on active frames in population raster plots of neural activity derived from two-photon calcium imaging in mice, or microelectrode array SUA in humans, as described above. In both mice and humans, temporal down-sampling was used so that in all datasets, one “frame” corresponded to 100ms of neural activity. Using the perplexity value with the lowest optimization error for each dataset (a value range from ~5-200 was tested), and an initial reduction to 25 dimensions of activity using principal component analysis (PCA), t-SNE was applied across 1000 repititions to produce a robust two-dimensional embedding space that could be conveniently visualized (Fig. 1G). This 2D embedding space of an entire experiment, in which every data point represents the activity of the entire recorded neural population per individual frame, was used for watershed segmentation in order to separate clusters of similar activity (Suppl. Fig. 1A). To this end, a density map was generated. A probability density function was calculated in the embedding space by convolving the embedded points with a Gaussian kernel; the standard deviation *σ* of the Gaussian was chosen to be 1/40 of the maximum value in the embedding space. To segment the density map, local maxima were identified in the density map, a binary map containing peak positions was generated, and peak points were dilated by three pixels. A distance map of the binary image was generated and inverted, and the peak positions were set to be minimum. Watershedding was performed on the inverted distance map, and the boundaries were defined with the resulting watershed segmentation. In this paper, a cluster identified by t-SNE and subsequent watershed segmentation, is called a “microstate”.

#### Lempel-Ziv-Welch Complexity analysis in mice and humans

The Lempel-Ziv Complexity (LZC), or sequence complexity, measures the complexity of a finite sequence of symbols by way of trying to compress the sequence (Ziv and Lempel, 1978). Here, the Lempel-Ziv-Welch (LZW) algorithm was used for this purpose, which is an encoding algorithm that works by creating a dictionary of common substrings (Welch, 1984). The Lempel-Ziv-Welch algorithm is an improved implementation over the initial algorithm proposed for calculating the LZC, in terms of computation cost. LZW was applied to raster firing matrices of recorded cellular calcium transients in mice or SUA in humans (Suppl. Fig. 2D). To arrive at its encoded form, first the binary matrix was transformed into a vector that was scanned by the LZW algorithm. A Matlab module was used to calculate LZWC [https://www.mathworks.com/matlabcentral/fileexchange/4899-lzw-compression-algorithm]. To make the results of the LZW algorithm comparable across experiments, and therefore independent of the number of imaged neurons per animal or single units recorded in the two human subjects, an upper and lower bound was established. This was done by running the LZW algorithm on a vector of the same size but with all zeros, as well as taking the average of random vectors of the same size. These were used to establish lower and upper bounds between complete order and randomness, respectively, and then normalize the length of the compressed vector of neural activity to arrive at a final value between 0 and 1, so that:

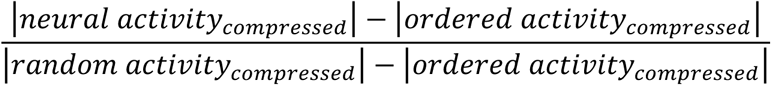

### Quantification and Statistical analysis

Unless stated otherwise, all values reported in this paper represent means ± s.e.m.. To determine statistical significance of differences between mean values measured under different experimental conditions in mice (n=7 animals, 5 conditions: awake, mild anesthesia, surgical anesthesia, deep anesthesia, and wake-up), 1-way anova was carried out (4 degrees of freedom), followed by a multiple comparison Bonferroni correction. Regarding statistical differences between observed and corresponding randomized numbers of unique microstates across experimental conditions in both mice and human subjects, observed single values were compared to a distribution of 100 values derived from randomized surrogate datasets. If the observed value was smaller/bigger than 95% of the randomized values, statistical significance was reached (p<0.05). P<0.01 was reached, if the observed value was smaller/bigger than all 100 randomized values. In this paper, statistical significance levels are depicted as * for p<0.05, ** for p<0.01, or *** for p<0.001.

### Additional Resources

MOCO (motion correction) is available on the Yuste lab website: http://www.columbia.edu/cu/biology/faculty/yuste/methods.html

### Resource Table

|  |  |  |
| --- | --- | --- |
| **Experimental Models: Mouse lines** |  |  |
| C57BL/6J-Tg(Thy1-GCaMP6f)GP5.11Dkim/J | Jackson Laboratory | RRID:IMSR_JAX:024276 |
| **Recording device in human patients** |  |  |
| Utah-style microelectrode array | Blackrock Microsystems |  |
| **Software and Algorithms** |  |  |
| FDA-approved neural signal processing system | Blackrock Microsystems |  |
| BIS Vista monitoring system | Aspect Medical Systems |  |
| ImageJ | https://imagej.nih.gov/ij/ |  |
| Moco | http://www.columbia.edu/cu/biology/faculty/yuste/methods.html |  |
| MATLAB(R2014b) MathWorks | MathWorks |  |
| Adobe Illustrator CS6 | Adobe |  |

Contributions: M.W. conceived of the project, designed and performed all experiments in mice, and wrote the manuscript. All authors discussed data, wrote the materials and methods section, and edited the manuscript. M.W. and S.H. pre-processed and analyzed data (calcium data, LFP, t-SNE, watershed segmentation, microstate analyses). E.H.S. processed and spike-sorted human data. E.P.H. independently discovered microstate alterations during LOC and provided LZW analysis for rodent and human data. B.G. and P.A.H. obtained and provided human data. R.Y. assembled and directed the team, supervised the study and provided guidance and funding.

## Acknowledgements

This work was supported by the NIMH (R01MH101218, R01MH100561) and NEI (DP1EY024503, R01EY011787). This material is also based upon work supported by, or in part by, the U. S. Army Research Laboratory and the U. S. Army Research Office under contract number W911NF-12-1-0594 (MURI). S.H. is a Howard Hughes Medical Institute International Student Research Fellow. We thank the Yuste lab members for input and constructive discussions.

This manuscript contains supplementary information (SI, Figures S1 to S3).

## References

Barth, A.L., and Poulet, J.F. (2012). Experimental evidence for sparse firing in the neocortex. Trends Neurosci 35, 345–355.

Barttfeld, P., Uhrig, L., Sitt, J.D., Sigman, M., Jarraya, B., and Dehaene, S. (2015). Signature of consciousness in the dynamics of resting-state brain activity. Proceedings of the National Academy of Sciences of the United States of America 112, 887–892.

Carrillo-Reid, L., Yang, W., Kang Miller, J.E., Peterka, D.S., and Yuste, R. (2017). Imaging and Optically Manipulating Neuronal Ensembles. Annu Rev Biophys.

Castaigne, P., Lhermitte, F., Buge, A., Escourolle, R., Hauw, J.J., and Lyon-Caen, O. (1981). Paramedian thalamic and midbrain infarct: clinical and neuropathological study. Ann Neurol 10, 127–148.

Cohen, M.R., and Kohn, A. (2011). Measuring and interpreting neuronal correlations. Nat Neurosci 14, 811–819.

Crick, F.C., and Koch, C. (2005). What is the function of the claustrum? Philos Trans R Soc Lond B Biol Sci 360, 1271–1279.

Dana, H., Chen, T.W., Hu, A., Shields, B.C., Guo, C., Looger, L.L., Kim, D.S., and Svoboda, K. (2014). Thy1-GCaMP6 transgenic mice for neuronal population imaging in vivo. PLoS One 9, e108697.

Flores, F.J., Hartnack, K.E., Fath, A.B., Kim, S.E., Wilson, M.A., Brown, E.N., and Purdon, P.L. (2017). Thalamocortical synchronization during induction and emergence from propofol-induced unconsciousness. Proc Natl Acad Sci U S A 114, E6660–E6668.

Herrera, C.G., Cadavieco, M.C., Jego, S., Ponomarenko, A., Korotkova, T., and Adamantidis, A. (2016). Hypothalamic feedforward inhibition of thalamocortical network controls arousal and consciousness. Nat Neurosci 19, 290–298.

Hopfield, J.J. (1982). Neural networks and physical systems with emergent collective computational abilities. Proceedings of the National Academy of Sciences of the United States of America 79, 2554–2558.

House, P.A., MacDonald, J.D., Tresco, P.A., and Normann, R.A. (2006). Acute microelectrode array implantation into human neocortex: preliminary technique and histological considerations. Neurosurg Focus 20, E4.

Hudetz, A.G., Liu, X., and Pillay, S. (2015). Dynamic repertoire of intrinsic brain states is reduced in propofol-induced unconsciousness. Brain Connect 5, 10–22.

Hudetz, A.G., Vizuete, J.A., Pillay, S., and Mashour, G.A. (2016). Repertoire of mesoscopic cortical activity is not reduced during anesthesia. Neuroscience 339, 402–417.

Ishizawa, Y., Ahmed, O.J., Patel, S.R., Gale, J.T., Sierra-Mercado, D., Brown, E.N., and Eskandar, E.N. (2016). Dynamics of Propofol-Induced Loss of Consciousness Across Primate Neocortex. J Neurosci 36, 7718–7726.

Jennett, B. (2002). The Vegetative State: Medical Facts, Ethical and Legal Dilemmas (Cambridge: Cambridge University Press).

Koch, C. (2012). In which I argue that consciousness is a fundamental property of complex things… (Cambridge, Massachusetts: MIT Press).

Lashley, K.S. (1929). Brain Mechanisms and Intelligence: A Quantitative Study of Injuries to the Brain (University of Chicago Press).

Lewis, L.D., Weiner, V.S., Mukamel, E.A., Donoghue, J.A., Eskandar, E.N., Madsen, J.R., Anderson, W.S., Hochberg, L.R., Cash, S.S., Brown, E.N., et al. (2012). Rapid fragmentation of neuronal networks at the onset of propofol-induced unconsciousness. Proc Natl Acad Sci U S A 109, E3377–3386.

Merker, B. (2007). Consciousness without a cerebral cortex: a challenge for neuroscience and medicine. Behav Brain Sci 30, 63–81; discussion 81-134.

Mhuircheartaigh, R.N., Rosenorn-Lanng, D., Wise, R., Jbabdi, S., Rogers, R., and Tracey, I. (2010). Cortical and subcortical connectivity changes during decreasing levels of consciousness in humans: a functional magnetic resonance imaging study using propofol. J Neurosci 30, 9095–9102.

Miller, J.E., Ayzenshtat, I., Carrillo-Reid, L., and Yuste, R. (2014a). Visual stimuli recruit intrinsically generated cortical ensembles. Proceedings of the National Academy of Sciences of the United States of America 111, E4053–4061.

Minert, A., Yatziv, S.L., and Devor, M. (2017). Location of the Mesopontine Neurons Responsible for Maintenance of Anesthetic Loss of Consciousness. J Neurosci 37, 9320–9331.

Moruzzi, G., and Magoun, H.W. (1949). Brain stem reticular formation and activation of the EEG. Electroencephalogr Clin Neurophysiol 1, 455–473.

Nyquist, H. (1924). Certain factors affecting telegraph speed. Bell Syst Tech J 3, 324–346.

Pavlov, P.I. (1927). Conditioned reflexes: An investigation of the physiological activity of the cerebral cortex (Oxford University Press).

Penfield, W.J., H. H. (1954). Epilepsy and the Functional Anatomy of the Human Brain (Little, Brown, Boston).

Pnevmatikakis, E.A., Soudry, D., Gao, Y., Machado, T.A., Merel, J., Pfau, D., Reardon, T., Mu, Y., Lacefield, C., Yang, W., et al. (2016). Simultaneous Denoising, Deconvolution, and Demixing of Calcium Imaging Data. Neuron 89, 285–299.

Schneider, C.A., Rasband, W.S., and Eliceiri, K.W. (2012). NIH Image to ImageJ: 25 years of image analysis. Nature methods 9, 671–675.

Schneidman, E., Berry, M.J., 2nd, Segev, R., and Bialek, W. (2006). Weak pairwise correlations imply strongly correlated network states in a neural population. Nature 440, 1007–1012.

Scoville, W.B., and Milner, B. (1957). Loss of recent memory after bilateral hippocampal lesions. J Neurol Neurosurg Psychiatry 20, 11–21.

Shoham, S., Fellows, M.R., and Normann, R.A. (2003). Robust, automatic spike sorting using mixtures of multivariate t-distributions. J Neurosci Methods 127, 111–122.

Steriade, M. (2003). The corticothalamic system in sleep. Front Biosci 8, d878–899.

Tononi, G. (2008). Consciousness as integrated information: a provisional manifesto. Biol Bull 215, 216–242.

Tononi, G., Boly, M., Massimini, M., and Koch, C. (2016). Integrated information theory: from consciousness to its physical substrate. Nat Rev Neurosci 17, 450–461.

van der Maaten, L.H. G. (2008). Visualizing Data using t-SNE. Journal of Machine Learning Research 9, 2579–2605.

Welch, T.A. (1984). A Technique for High-Performance Data-Compression. Computer 17, 8–19.

Yang, W., Miller, J.E., Carrillo-Reid, L., Pnevmatikakis, E., Paninski, L., Yuste, R., and Peterka, D.S. (2016). Simultaneous Multi-plane Imaging of Neural Circuits. Neuron 89, 269–284.

Yang, W., and Yuste, R. (2017). In vivo imaging of neural activity. Nat Methods 14, 349–359.

Yuste, R., and Denk, W. (1995). Dendritic spines as basic units of synaptic integration. Nature 375, 682–684.

Yuste, R., and Katz, L.C. (1991). Control of postsynaptic Ca2+ influx in developing neocortex by excitatory and inhibitory neurotransmitters. Neuron 6, 333–344.

Ziv, J., and Lempel, A. (1978). Compression of Individual Sequences Via Variable-Rate Coding. Ieee T Inform Theory 24, 530–536.

